# A Universal Duplex Sequencing Approach for Accurate Detection of Somatic Mutations

**DOI:** 10.1101/2025.09.14.676103

**Authors:** Shuvro P. Nandi, Yuhe Cheng, Shams Al-Azzam, Safa Saeed, Audrey Kristin, Nadia Sunico, Isabella R. Stuewe, Zichen Jiang, Luka Culibrk, Maria Zhivagui, Xiaoxu Yang, Rachel M. Wise, Foster C Jacobs, Bérénice Chavanel, Michael Korenjak, Mia Petljak, Silvia Balbo, Laurie G. Hudson, Ke Jian Liu, Jiri Zavadil, Joseph G. Gleeson, Ludmil B Alexandrov

**Author notes:** These authors contributed equally (Shuvro P. Nandi, Yuhe Cheng).

## Abstract

Ultra-accurate detection of rare somatic mutations is critical for understanding mutational processes in human disease, aging, and environmental exposures, yet current methods are limited by error rates, restricted genome coverage, and high DNA input. We present UDSeq, a duplex sequencing protocol combining random fragmentation, efficient UMI ligation, and quantitative input control to achieve near-complete genome/exome representation from as little as 100 pg DNA. Benchmarking in human sperm estimates a UDSeq error rate of ∼2.5×10^-9^ per base pair. UDSeq captures mutational signatures from heterogeneous populations without clonal expansion, reproduces exposure-specific patterns in cell lines and rodent models, and enables cross-species profiling. Compared with prior duplex methods, UDSeq yields up to fourfold more usable duplex molecules, improves library conversion, and remains cost-effective. We include a step-by-step protocol with quality-control checkpoints for fragment size, ligation yield, library conversion, and duplication rate. UDSeq provides a scalable, low-input platform for accurate profiling of somatic mutagenesis.

## INTRODUCTION

Somatic mutations, which can be caused by both endogenous and exogenous mutagenic processes, are present in all human cells^1^. These mutations accumulate gradually over time and often go unnoticed, as most have minimal or no effect on cellular function^1–3^. However, certain mutations can disrupt key biological processes^4^, lead to cell death^5^, or confer a selective growth advantage, resulting in clonal expansion^6,7^. Cancer is the most well-known example of a disease driven by somatic mutations, where specific alterations initiate tumorigenesis^8^, promote progression^9^, and can confer treatment resistance^10^. Beyond cancer, somatic mutations are increasingly recognized as contributors to other diseases^11^, including neurodegenerative disorders^12^ and cardiovascular conditions^13^. However, their roles in these areas remain much less studied, as detecting low-frequency mutations in non-cancer tissues presents significant technical challenges^14,15^.

From a technical standpoint, the study of somatic mutations in cancer has been greatly facilitated by the fact that most tumors originate from a single mutated cell, whose proliferation produces thousands of descendants carrying the same mutations as the progenitor^8^. Although additional mutations may emerge during clonal expansion, the original mutations remain uniformly present across the tumor tissue^9^. This clonal nature enables reliable detection of tumor mutations with conventional sequencing, despite its inherent error rates ranging from one error per 1,000 base pairs (bp; *i.e.,* 10^-3^ per bp) to six errors per 1,000 bp (*i.e.,* 6 × 10^-3^ per bp)^16^. Because these shared mutations appear in every cancer cell, repeated resequencing generates a strong, reproducible signal that stands out from the random noise of sequencing errors. In contrast, detecting somatic mutations in most non-cancer tissues is far more challenging, as each cell typically harbors a unique set of mutations^17,18^. As such, accurately studying somatic mutations in non-cancerous healthy or diseased tissues requires methods with error rates below 1 × 10^-8^ errors per bp (less than one error per 100 million sequenced bp)^18^.

The need for sequencing protocols with low error rates is further evident in efforts to evaluate the mutagenic potential of known carcinogens in experimental systems. For example, *in vitro* studies have required intricate experimental setups, where cells are exposed to a potential mutagenic carcinogen, followed by isolation and clonal expansion of single cells from the exposed population before sequencing^19,20^. The clonal amplification step, though labor-intensive, is crucial for accurately detecting mutations, as it ensures that mutations present in the progeny of a single cell can be distinguished from background noise^19,20^. Without this step, bulk sequencing cannot distinguish low-frequency mutations from background noise, since each cell harbors a distinct set of somatic mutations.

To overcome these limitations, several approaches have been developed to detect rare somatic mutations. Single-cell DNA sequencing^21–25^ and single-cell clonal expansion^26–28^ can, in principle, resolve mutations at the level of individual cells^26,28^. However, these methods are labor-intensive, costly, and require sequencing large numbers of cells to obtain a representative view of tissue-wide mutagenesis. Additionally, amplification artifacts^29^ and allelic dropout^30^ can compromise accuracy for detecting rare mutations. As a scalable alternative, duplex sequencing has emerged as a powerful method for profiling somatic mutations at extremely low frequencies^31^. By exploiting DNA’s double-stranded nature, duplex sequencing independently evaluates each strand^32,33^ and confirms mutations only when they are independently detected on both strands, greatly reducing error rates and enabling confident identification of rare somatic mutations.

Advances in duplex sequencing have improved the detection of rare somatic mutations, but no current method offers universal, single-molecule resolution in a cost-effective format that supports both whole-genome profiling and targeted capture from limited input DNA across species. An ideal approach would achieve an error rate below 10^-8^, offer full genome compatibility for profiling human tissues, human and non-human model systems, and non-model organisms, while requiring minimal DNA input, supporting targeted enrichment, and remaining cost-effective. To address this need, we developed Universal Duplex Sequencing (UDSeq), a novel single-molecule duplex sequencing protocol for rapid, accurate detection of rare somatic mutations across diverse biological systems. We put UDSeq in the context of existing error-corrected sequencing methods, demonstrating superior performance and broad applicability. To showcase its versatility, we applied UDSeq to samples from humans, mice, rats, chickens, and sheep, successfully detecting mutations in whole genomes and targeted regions derived from cell lines and multiple tissue types.

## RESULTS

### Overview of Existing Error-Corrected Sequencing Methods

Over the past decade, duplex sequencing has revolutionized the detection of rare somatic mutations^18^. In this approach, a ‘duplex consensus’ is generated by independently sequencing both the Watson and Crick strands of the same DNA molecule and typically confirming mutations only if present on both strands. Most protocols use unique molecular identifiers (UMIs) and exploit DNA strand complementarity to perform this process^33^. To our knowledge, the first method of this kind was introduced in 2012, enabling sequencing of small genomic panels—generally under 1 megabase—with error rates of approximately 10^-7^ errors per bp^32^. However, this original DupSeq method had low efficiency in generating duplex consensuses^32^, limiting its practical application by necessitating large amounts of input DNA and extensive sequencing. Subsequently, *Hoang et al.* developed BotSeqS to address this challenge, introducing a dilution step immediately before library amplification^34^. This dilution step creates a bottleneck, enabling efficient random sampling of double-stranded template molecules and substantially reducing the required amount of sequencing. Notably, BotSeqS could be applied to input DNA amounts as low as 50 nanogram (ng). Despite this improvement, the error rate of BotSeqS was similar to that of DupSeq, with independent analysis estimating it at ∼2 × 10^−7^ errors per bp^35^.

To further enhance BotSeqS and reduce error rates, NanoSeqV1 was developed by incorporating optimized DNA fragmentation and restrictive end repair method during library preparation by replacing sonication and end repair with restriction enzyme-based fragmentation using HpyCH4V^35^. This innovation reduced the error rate to approximately 5 × 10^-9^ errors per base pair and allowed NanoSeqV1 to be applied to input DNA amounts as low as 50 ng. Nonetheless, it limited coverage to only 30% of the human genome and restricted its applicability to other genomes due to the specificity of the restriction enzyme^35^. More recently, the CODEC (Concatenating Original Duplex for Error Correction) method employed specially designed quadruplex adaptors to physically link the Watson and Crick strands into a single-duplex molecule, enabling sequencing on a standard Illumina short-read platform^36^. The original CODEC method achieved error rates of approximately 10^-7^ errors per bp, comparable to those of DupSeq and BotSeqS, while offering greater cost-effectiveness compared to DupSeq^36^. Additionally, the original CODEC method could be applied to input DNA amounts as low as 2.5 ng. A modified version of the CODEC protocol, incorporating fragmentation steps similar to those used by NanoSeqV1, reduced the error rate to approximately 10^-8^ errors per bp. However, this modified version inherited the same limitations as NanoSeqV1, including coverage restricted to only 30% of the human genome^36^. Additionally, to overcome the partial genome coverage limitations of the original NanoSeq, a second version—NanoSeqV2— was developed using an alternative genome fragmentation strategy. Nonetheless, its low library conversion efficiency necessitates a substantially larger amount of input DNA, which may restrict its broader applicability^37^.

The previously described methods relied entirely on short-read sequencing based on duplex sequencing. In contrast, a recently developed long-read sequencing technique, HiDEF-seq (Hairpin Duplex Enhanced Fidelity sequencing), leveraged the PacBio platform to achieve single-molecule fidelity^38^. HiDEF-seq utilized the inherent single-molecule nature of PacBio’s technology^39^ to achieve high accuracy, performing 5 to 20 sequencing passes per strand with estimated error rates below 10^-9^ errors per bp^38^. Notably, HiDEF-seq can resolve some single-strand mismatches, a capability not achievable with other duplex sequencing methods. However, it required a high input of DNA and incurred higher costs due to the expense of PacBio long-read sequencing compared to short-read technologies. Specifically, it needed at least 500 ng of high-quality DNA or 1,500 ng of degraded DNA to achieve 40% genome coverage, with even larger amounts required for complete genome sequencing^38^.

Each of the previously discussed approaches—DupSeq, BotSeqS, NanoSeq, CODEC, and HiDEF-seq—represents a significant advancement in the detection of rare somatic mutations, introducing innovative methodologies, enhanced efficiency, and reduced error rates (**Table 1**). However, each method also comes with its own set of limitations, including challenges related to efficiency, error rates, genome coverage, DNA input requirements, or cost. To overcome many of these limitations, we present UDSeq, a single-molecule duplex sequencing protocol optimized for rapid, accurate detection of rare somatic mutations (**Table 1**), with the complete protocol provided in **Supplementary Note 1**.

**Table 1:** Comparative overviews of DupSeq, BothSeq, NanoSeq, HiDEF-seq, CODEC, and UDSeq.

| Sequencing protocol | Primary innovation | Genome coverage | Minimum input DNA | Approx. error rate (per bp) | Key limitations |
| --- | --- | --- | --- | --- | --- |
| **DupSeq**<sup>32</sup> | Independent tagging and sequencing of both DNA strands to enable duplex consensus mutation calling | Targeted panels | ~1 µg | $2 \times 10^{-7}$ | Requires high DNA input; generally limited to defined panels; labor-intensive workflow; High error rate when compared to other protocols |
| **BotSeqS**<sup>34</sup> | Bottleneck dilution to enrich randomly sampled duplex DNA molecules for consensus mutation calling | Genome-wide or targeted | 50 ng | $2 \times 10^{-7}$ | High error rate when compared to other protocols. Low efficiency. |
| **NanoSeqV1**<sup>35</sup> | Removes end-repair-associated errors via restriction-enzyme fragmentation | ~30% of genome | 50 ng | $5 \times 10^{-9}$ | Restriction sites limit coverage and species portability; needs redesign for new targets |
| **NanoSeqV2**<sup>37</sup> | Alternative genome fragmentation methods that provide full genome coverage whilst retaining the original error rates | Genome-wide or targeted | 30 ng | $5 \times 10^{-9}$ | Low efficiency. Still unpublished but available as preprint |
| **HiDEF-seq**<sup>38</sup> | High-fidelity duplex consensus from PacBio long-read sequencing with detection of single-strand mismatches | ~40% of genome | 500 ng – 1.5 µg | $4 \times 10^{-9}$ | Complex workflow, elevated sequencing depth required; partial genome coverage; high cost |
| **CODEC**<sup>36</sup> | Custom quadruplex adapters physically linking Watson & Crick strands | Genome-wide or targeted | 2.5 ng | $1 \times 10^{-7}$ | Requires specialized reagents, sequencing customization, and intensive library preparation; higher error rate than other protocols. |
| **CODEC-HpyCH4V**<sup>36</sup> | Restriction enzyme–based CODEC enabling targeted duplex capture | ~30% of genome | 50 ng | $3 \times 10^{-8}$ | Requires specialized reagents, sequencing customization, and intensive library preparation; higher error rate than other protocols. |
| **UDSeq** | Near-complete genome coverage with ultra-low input via random fragmentation and efficient duplex consensus. | Genome-wide or targeted | 100 pg | $2.5 \times 10^{-9}$ | Similar to all other duplex methods, does not capture large structural variants or copy-number changes |

### Innovation Over Prior Protocols

To develop the UDSeq protocol, we built upon the advances of NanoSeqV1 over BotSeqS and introduced targeted innovations that overcome key limitations in existing duplex sequencing methods. Each improvement in the protocol was designed not only to enhance performance but also to expand the scope, versatility, and practicality of single-molecule mutation detection (**Figure 1*a***; **Supplementary Figure 1a**).

**Figure 1:**
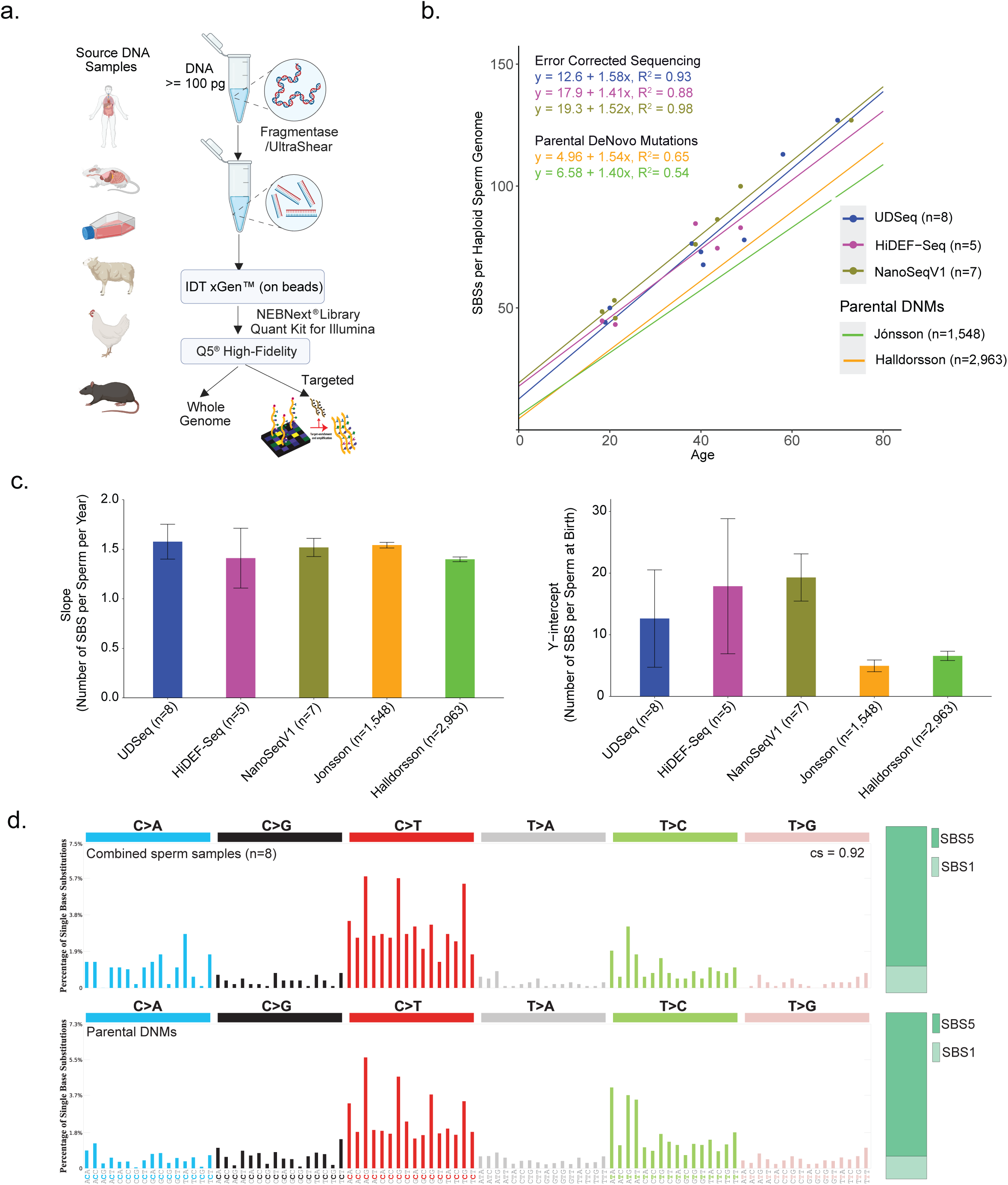
**Overview and validation of UDSeq for accurate detection of somatic mutations. *(a)*** High-level workflow illustrating the versatility of UDSeq across whole-genome and targeted sequencing approaches. ***(b)*** Comparison of error rates among UDSeq, other duplex sequencing methods, and germline *de novo* mutation (DNM) studies. UDSeq and other duplex approaches were applied to human sperm samples, whereas the DNM studies analyzed germline data from trios. Scatter plot shows the relationship between paternal age and the number of single base substitutions (SBSs) per haploid sperm genome across different sequencing approaches. Data points represent individual samples analyzed by UDSeq (*n*=8), HiDEF-Seq (*n*=5), and NanoSeqV1 (*n*=7), as well as parental DNM estimates from Jónsson (*n*=1,548) and Halldorsson (*n*=2,963). Regression lines with corresponding equations and coefficients of determination (R²) are shown for each dataset: error-corrected sequencing (UDSeq, HiDEF-Seq, NanoSeqV1) and parental DNM studies. ***(c)*** Left panel shows estimated slopes for the number of SBS accumulated per haploid sperm genome per year, and the right panel shows estimated y-intercepts representing the predicted number of SBS present at birth. Both values are derived from the regression analyses in panel b. Error bars indicate standard error of the estimate. ***(d)*** Top panel shows the SBS-96 mutational profile from sperm samples analyzed by UDSeq (*n*=8), and the bottom panel shows the SBS-96 profile from parental DNMs. The SBS-96 profile encompasses all single-base substitutions (C>A, C>G, C>T, T>A, T>C, and T>G) and their immediate trinucleotide sequence context. The two profiles have a cosine similarity of 0.92. Relative contributions of the aging-associated signatures SBS1 and SBS5 are shown on the right for each profile.

First, we replaced sonication-and HpyCH4V-based fragmentation with random fragmentation using either NEBNext dsDNA Fragmentase (M0348L) or UltraShear (M7634L). Both methods enable unbiased fragmentation across the genome, allowing UDSeq to achieve near-complete coverage (≥95%; comparable to bulk sequencing) of the genome and exome—an advance over NanoSeqV1, which was limited to ∼30% of the genome (**Supplementary Figure 1b**), and comparable to NanoSeqV2, which also provides near-complete coverage^37^. dsDNA Fragmentase is more cost-effective and widely accessible, though it produces short overhangs that require additional trimming during bioinformatics processing. In contrast, UltraShear generates highly uniform fragment sizes without overhangs, reducing computational preprocessing steps—but at higher reagent cost. This flexibility allows users to balance performance, cost, and bioinformatics complexity based on experimental needs.

Second, we adopted the xGen™ cfDNA & FFPE DNA Kit (IDT) for ligation of unique UMIs. This kit provides high ligation efficiency even with minimal DNA input, enabling accurate duplex sequencing from as little as 0.1 ng (100 picograms) of starting material. Compared to NanoSeqV2, it delivers a substantial improvement in library conversion efficiency— yielding up to four times more femtomoles of usable library from the same input DNA (p=0.00022; **Supplementary Figure 1c**). This enhancement reduces input requirements and broadens applicability to samples with limited DNA, such as clinical biopsies or environmental isolates.

Third, we incorporated accurate quantification of UMI-ligated molecules using the NEBNext Library Quant Kit for Illumina and iTaq Universal SYBR Green Supermix on a Bio-Rad real-time PCR system. This ensures precise input into PCR amplification, reducing over-amplification artifacts and preserving single-molecule fidelity. PCR was then performed with UDI primers (IDT) and NEBNext® Ultra™ II Q5® Master Mix (M0544L), which maintains high fidelity during amplification.

Finally, by integrating random fragmentation with low-input, high-efficiency UMI ligation and precise quantification, UDSeq uniquely enables ultra-accurate, single-molecule somatic mutation detection across species, with support for whole-genome coverage or targeted panels—even from limited input material. Despite offering substantial advantages in sensitivity, flexibility, and scalability, the protocol remains cost-efficient— comparable to or even lower in cost than other duplex sequencing methods. (**Supplementary Figure 1d**). In the sections that follow, we systematically evaluate the protocol’s error rate and demonstrate its applicability across a range of experimental settings. Together, these advances position UDSeq as a cost-effective, scalable platform for widespread genomic applications (**Figure 1a**; **Supplementary Figure 1a).**

### Assessing and Comparing the Error Rate of UDSeq

To assess the error rate of UDSeq, we sequenced DNA extracted from sperm samples provided by eight males ranging in age from 19 to 70 years. We compared our results with two large-scale Icelandic population studies of trios (mother, father, and child), which estimated sperm mutation rates in fathers at different ages based on phasing *de novo* mutations (DNM) observed in the offspring^40,41^. These DNM trio studies reported sperm mutation rates of 1.54 and 1.40 single base substitutions (SBS) per year, respectively, with the number of SBS at birth (i.e., age zero) estimated at 4.96 and 6.58, respectively (**Figure 1*b-c***). Consistent with the estimates from the DNM trio studies, the UDSeq data revealed that sperm accumulate 1.58 SBSs per year and an estimated number of mutations at age zero of 12.60. By analyzing the difference in mutation rates at age zero between UDSeq and the DNM trio studies, we estimated that the error rate of UDSeq is between 6 and 7.6 artifactual SBS per sequenced haploid sperm sample containing approximately three billion base pairs. This corresponds to an error rate of approximately 2.5 × 10^-9^ errors per bp (**Supplementary Figure 1d**). Applying the same approach to previously sequenced sperm samples, we also estimated the error rates of NanoSeqV1 (*n*=7 sperm samples) and HiDEF-seq (*n*=5) which yielded error rates of about 4.8 × 10^-9^ and 4.3 × 10^-9^ errors per bp, respectively (**Figure 1*c***). Although derived using a different approach, our estimated error rate closely matches values reported in previous studies— for example, 4.8 × 10^-9^ in our analysis compared to 5 × 10^-9^ for NanoSeqV1 in their original publication^35^. Overall, given the sample sizes of sperm samples, the error rates of UDSeq, NanoSeq, and HiDEF-seq were effectively similar, with less than 5 mutations per billion sequenced base pairs (*i.e.*, <5 × 10^-9^ errors per bp; **Figure 1*c***). Lastly, as expected^42,43^, the mutational patterns observed in the eight sperm samples profiled by UDSeq exhibited the patterns of clock-like signatures SBS1 and SBS5, closely resembled that of paternal *de novo* mutations^41^ (cosine similarity = 0.92; **Figure 1*d***).

### *In vitro* Assessment of Mutagenesis

Traditional sequencing protocols for evaluating environmental carcinogen exposure *in vitro* generally require months of precise exposure, clonal expansion, and sequencing^19,44,45^ (**Figure 2*a***), whereas UDSeq enables direct mutational detection in heterogeneous cell populations, significantly reducing the timeline (**Figure 2*b***). To showcase the utility of the UDSeq protocol, we applied it to three human cell lines exposed to four environmental carcinogens with well-documented mutagenic properties. These *in vitro* experiments included: *(i)* HepG2 human liver cancer cell line, derived from a well-differentiated hepatocellular carcinoma, exposed to 4-Nitroquinoline 1-oxide (4NQO; 0.5 µM for 4 hours) and aristolochic acid-I (AA-I; 80µM for 24 hours); *(ii)* immortalized normal oral keratinocytes (NOK) exposed to the tobacco specific nitrosamine 4-methylnitrosamino-1-(3-pyridyl)-1-butanone (NNK)^46^; and *(iii)* N/TERT-1 keratinocytes exposed to solar-simulated ultraviolet-light radiation (ssUVR; 3 KJ/m^2^)^47^. A heterogeneous population of cells was cultured for a single passage following exposure, after which DNA was extracted and subjected to UDSeq at whole-genome resolution (**Figure 2*b***). The resulting mutational profiles from this duplex sequencing approach were then compared to those obtained from clonally expanded cells that were exposed to the same carcinogens (**Figure 2*b***).

**Figure 2:**
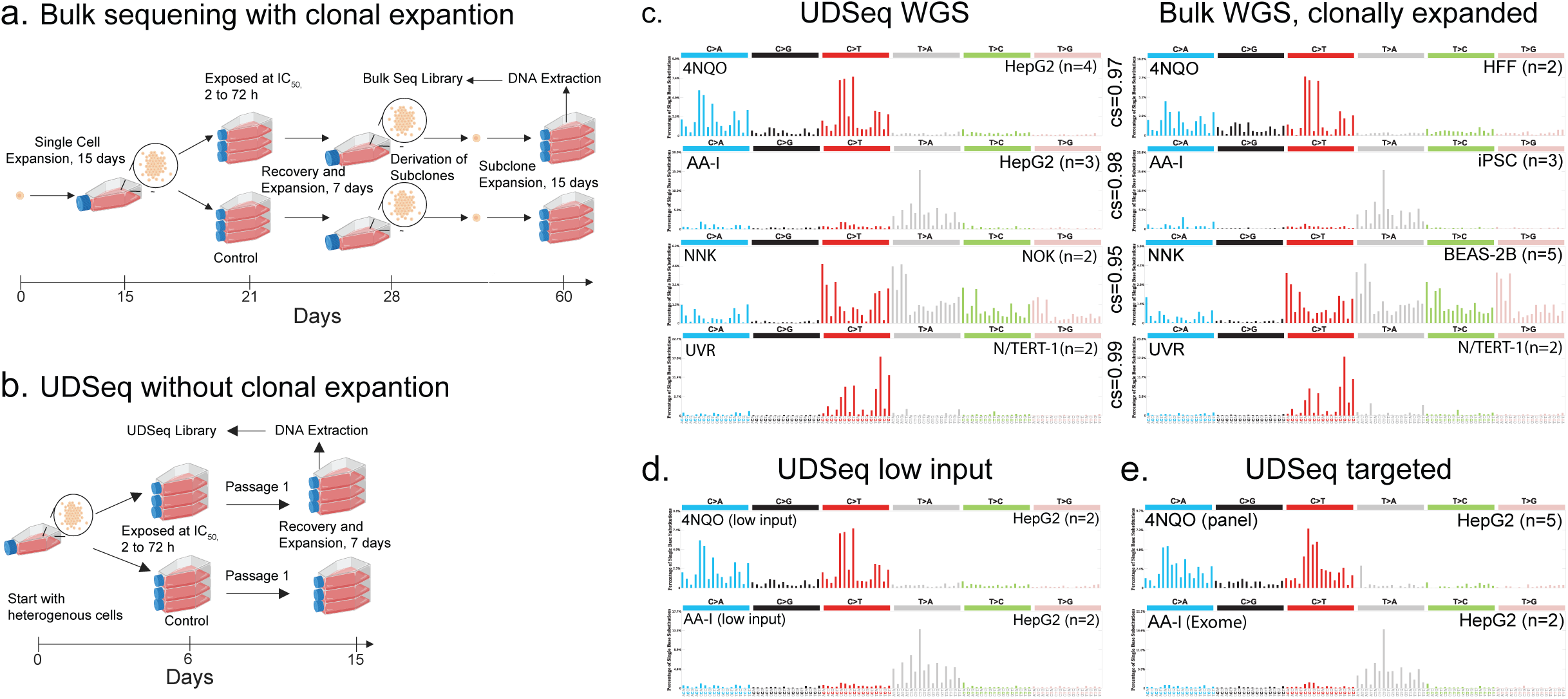
**UDSeq enables rapid, ultra-low-input, and versatile assessment of *in vitro* mutational profiles. *(a)*** Schematic workflow for *in vitro* mutagenesis assessment using single-cell clonal expansion followed by bulk sequencing, a process requiring 60 days or more. ***(b)*** Alternative workflow using UDSeq without clonal expansion, enabling mutagenesis assessment in as little as 15 days. ***(c)*** SBS-96 mutational profiles of environmental mutagens. Each SBS-96 profile represents all single-base substitutions (C>A, C>G, C>T, T>A, T>C, T>G) within their trinucleotide sequence context. *Left:* UDSeq-derived SBS-96 mutational profiles from human cell lines exposed to mutagens. *Right:* SBS-96 derived mutational profiles from human cell lines exposed to mutagens using single-cell clonal expansion followed by bulk sequencing. Cosine similarities (cs) between UDSeq-derived and bulk sequencing–derived profiles are shown between the panels. Mutagen names are indicated in the top left corner, and the corresponding cell line with replicate number is shown in the top right corner of each panel. ***(d)*** SBS-96 mutational profiles generated from as little as 100 picograms of DNA using UDSeq. SBS-96 mutational profiles from environmental mutagen exposures are shown with cell line names and replicate numbers indicated as in *(c)*. ***(e)*** Demonstration of UDSeq’s versatility across exome and targeted sequencing. SBS-96 mutational profiles from environmental mutagen exposures are shown for both exome and targeted panels, with cell line names and replicate numbers indicated as in *(c)*. Abbreviations: 4NQO, 4-Nitroquinoline 1-oxide; AA-I, aristolochic acid I; BEAS-2B, immortalized human bronchial epithelial cell line; HFF, human foreskin fibroblast; HepG2, human liver cancer cell line derived from hepatocellular carcinoma; IC50, half-maximal inhibitory concentration; iPSC, induced pluripotent stem cells; NNK, 4-(methylnitrosamino)-1-(3-pyridyl)-1-butanone; NOK, oral and epidermal keratinocytes; UVR, ultraviolet radiation.

The mutational profile induced by AA-I closely matched the COSMIC signature SBS22a, consistent with SBS22a’s AA-I etiology as established from human cancer samples^48^ and *in vitro* models^49^. The profile also showed strong concordance with bulk sequencing data derived from clonally expanded AA-I-exposed cells (cosine similarity = 0.98; **Figure 2*c***). Similarly, the pattern of ssUVR matched the known ssUVR-light associated COSMIC experimental mutational signature for solar simulated radiation^47^, while the pattern of 4NQO was identical to that observed in clonally expanded human cells exposed to 4NQO (**Figure 2*c***). Lastly, for NNK acetate, the mutational profiles were also nearly identical to the one found in clonally expanded human lung cells exposed to the same compound^50^. Overall, these results confirm that UDSeq can replicate mutational patterns observed in clonally expanded cells, without the time or resource burden of extended culture.

The previously generated *in vitro* results utilized 100 ng of input DNA (**Figure 2*c***). To evaluate UDSeq’s performance with low-input DNA, we applied it to 100 picograms (pg) of DNA 4NQO-and AA-I-exposed HepG2 cells. Across all conditions, UDSeq reliably detected the expected mutational patterns, yielding results consistent with those obtained from higher DNA input (**Figure 2*d***).

To demonstrate UDSeq’s versatility in generating custom pull-down sequencing, we also performed whole-exome sequencing (using xGen™ Exome Hybridization Panel) and targeted gene panel sequencing encompassing 127 known cancer-associated genes (xGen™ Pan-Cancer Hybridization Panel). The resulting mutational profiles closely matched expectations, with AA-I exposure aligning with SBS22a (cosine similarity = 0.98) and 4NQO exposure mirroring patterns observed in HepG2 cells (cosine similarity = 0.94; **Figure 2*e***).

### *In vivo* Assessment of Mutagenesis

To showcase the utility of the UDSeq protocol for assessing *in vivo* mutagenesis, we exposed SKH-1 hairless mice to ssUVR (14 kJ/m^2^; ∼0.5 minimal erythema dose) three times per week for 30 weeks (**Figure 3*a***) and F344 rats to 5 parts per million NNK in drinking water for 15 weeks (**Figure 3*b***), and compared their mutational profiles to those of unexposed controls. As expected, in SKH-1 hairless mice, mutational burden analysis revealed 105-fold and 6.5-fold higher mutational loads in the dorsal and ventral skin of ssUVR-exposed mice, respectively (p<0.05), compared to controls (**Figure 3*a***). ssUVR-associated mutational signatures were present in the dorsal and ventral skin of exposed mice but absent in the skin of unexposed controls (**Figure 3*a***). Similarly, F344 rats exposed to NNK exhibited a 4.7-fold higher mutational burden compared to controls (p=0.013; **Figure 3*b***), with an NNK-specific mutational pattern closely resembling that observed in cell line experiments (**Figure 2*b***; cosine similarity = 0.90).

**Figure 3:**
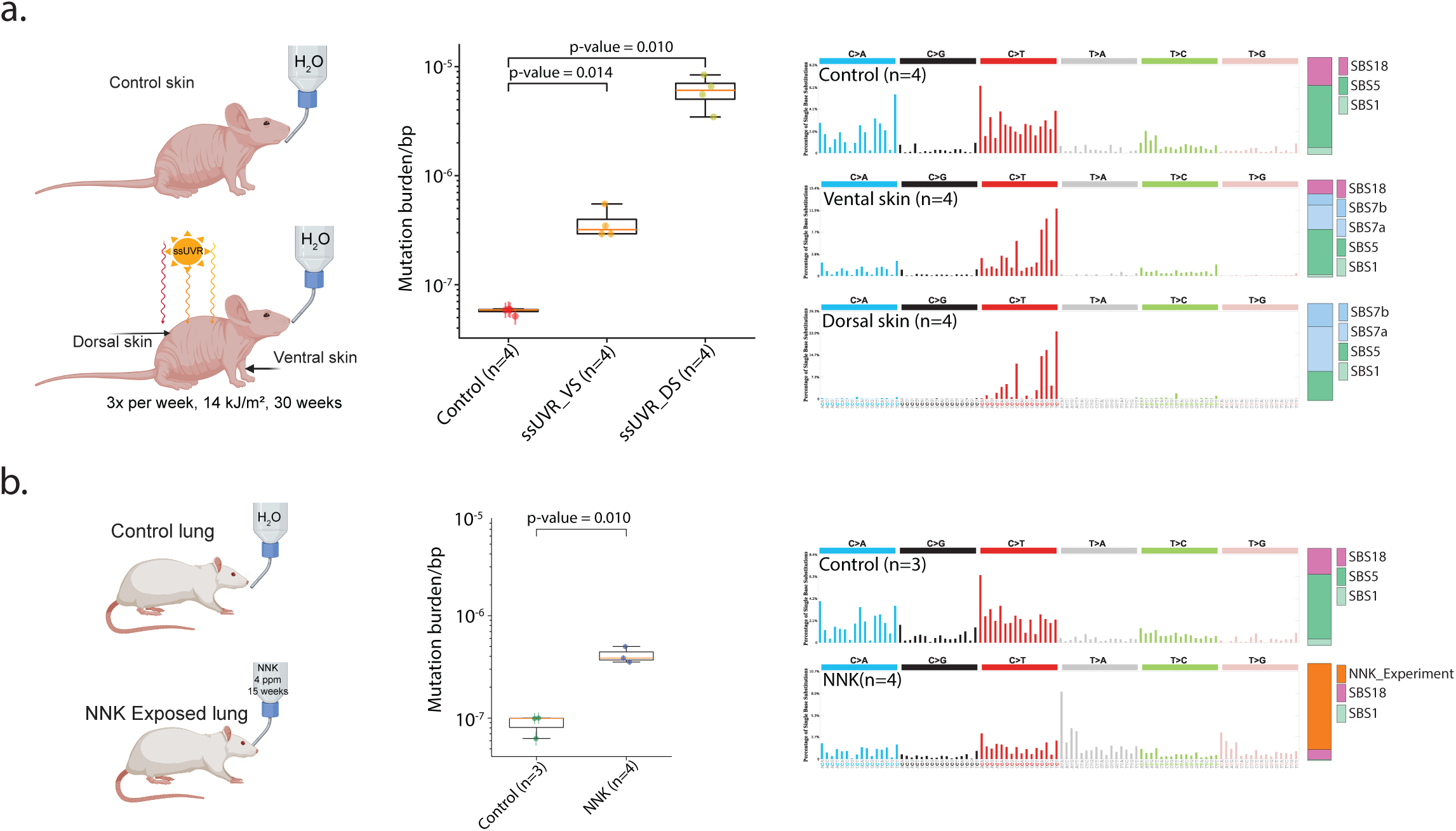
**UDSeq-based *in vivo* mutagenesis in mouse and rat models. *(a)*** *Left:* Schematic of the in vivo mutagenesis workflow in SKH-1 hairless mice, with cohorts either unexposed (controls) or subjected to solar-simulated UVR (14 kJ/m²; ∼0.5 minimal erythema dose) three times per week for 30 weeks. *Middle:* Box plots showing mutation burden per base pair in ventral skin from control mice (Control; *n*=4), ventral skin from ssUVR-exposed mice (ssUVR_VS, *n*=4), and dorsal skin from ssUVR-exposed mice (ssUVR_DS, *n*=4). The y-axis represents mutation burden per base pair on a log scale. Horizontal lines within boxes indicate medians; boxes represent interquartile ranges (IQR), and whiskers extend to 1.5× IQR. P-values were calculated using a two-sided t-test: control vs. ssUVR_VS, p=0.014; ssUVR_VS vs. ssUVR_DS, p=0.010. *Right*: SBS-96 mutational profiles of control, ssUVR_VS, and ssUVR_DS skin samples, with contributing COSMIC reference mutational signatures shown adjacent to each profile. Each SBS-96 profile represents all single-base substitutions (C>A, C>G, C>T, T>A, T>C, T>G) within their trinucleotide sequence context. ***(b)*** *Left:* Schematic of the in vivo mutagenesis workflow in F344 rats: control and 4-(methylnitrosamino)-1-(3-pyridyl)-1-butanone (NNK)–exposed groups (NNK administered in drinking water for 15 weeks), with lung tissue collected for analysis. *Middle:* Box plots showing mutation burden per base pair in lung tissue for control (*n*=3) and NNK-exposed (*n*=4) lungs. The y-axis represents mutation burden per base pair on a log scale, and the format of the box plots is identical to the one in *(b)*. P-values were calculated using a two-sided t-test: control vs. NNK, p=0.010. *Right*: SBS-96 mutational profiles of control and NNK-exposed lung tissues, with contributing COSMIC reference signatures shown alongside each profile. The *in vitro*–derived NNK experimental signature from *Figure 2c* was included in the assignment and was detected exclusively in the NNK-exposed samples.

Additionally, we evaluated the capability of UDSeq to profile tissues from non-model organisms by whole-genome sequencing breast, pancreas, and skin tissues from healthy chickens, as well as different layers of kidney tissue samples from healthy sheep. As anticipated^51^, the tissues from both chickens and sheep displayed distinct patterns of clock-like mutational signatures SBS1 and SBS5, along with SBS18 in skin tissue of chickens (**Supplementary Figure 3a-c**). Furthermore, we observed that the cortex has a 1.49-fold higher mutational burden than the medulla in the same kidney samples (**Supplementary Figure 3c**)^52^.

### Examining Mutational Processes in Healthy Human Tissues

To demonstrate UDSeq’s ability to study mutational processes in normal somatic tissues of healthy individuals, we applied the protocol at whole-genome resolution to five organs from a single 70-year-old individual: left cortex, right cortex, left kidney, right kidney, and liver (**Figure 4*a***). The data revealed that the brain had the lowest mutational burden, followed by the kidneys and liver (**Figure 4*a***). Interestingly, the left kidney exhibited more mutations than the right kidney. However, since DNA was extracted from bulk tissue, this difference may stem from variations in capturing the kidney cortex and medulla, as observed in prior reports^52^ and our data from sheep kidney (**Supplementary Figure 3c**). To investigate the mutational processes active in these organs, we analyzed the mutational signatures present^53^ across tissues and identified patterns consistent with known biology. As expected, clock-like mutational signatures SBS1 and SBS5— associated with cell proliferation and aging—were detected in all samples (**Figure 4*b***). SBS40, a signature commonly observed in renal tissues and cancers despite its unknown etiology^54^, was present in both kidney samples. SBS4, which is associated with tobacco smoking across multiple cancer types and has also been observed in liver cancer^55^, was uniquely detected in the liver, although the smoking status of the donor was unavailable. Together, these findings confirm the ability to detect the presence of expected tissue-specific mutational processes (**Figure 4*b***).

**Figure 4:**
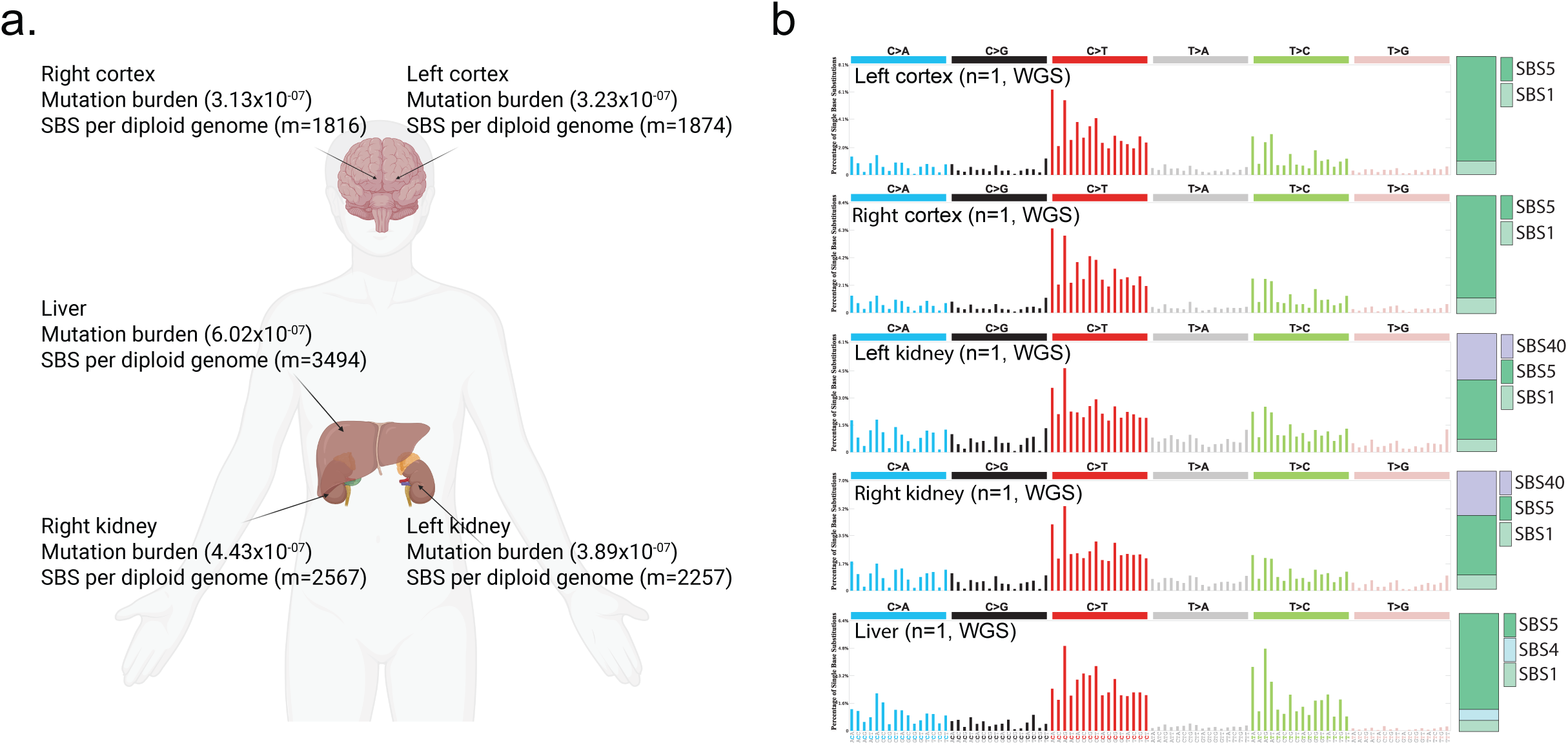
**UDSeq-based mutational burden and mutational profiles across healthy human tissues. *(a)*** Schematic of the human body highlighting sampled organs and their mutation burden estimates from a single 70-year-old individual. Mutation burden is expressed as single base substitutions (SBS) per base pair, and the total number of SBS per diploid genome is shown for each organ (denoted by *m*). ***(b)*** SBS-96 mutational profiles are shown for each tissue. Each SBS-96 profile represents all single-base substitutions (C>A, C>G, C>T, T>A, T>C, T>G) within their trinucleotide sequence context. Contributing COSMIC mutational signatures are displayed adjacent to each profile. Y-axis scales are adjusted individually to optimally display the percentage of mutations within each tissue.

## DISCUSSION

In this study, we introduce UDSeq, a novel and cost-efficient single-molecule duplex sequencing protocol designed to overcome the key limitations of existing error-corrected sequencing technologies. By enabling both whole-genome and targeted sequencing from as little as 100 picograms of input DNA, UDSeq combines ultra-low error rates (∼2.5 × 10^-9^ errors per base pair; **Table 1**) with high sensitivity and versatility, making it well-suited for detecting rare somatic mutations across a wide range of biological contexts. Benchmarking against human sperm DNA validated its accuracy, yielding mutation rates that align with parent-offspring trio-based *de novo* mutation studies and faithfully recapitulating clock-like mutational signatures SBS1 and SBS5.

Unlike prior duplex sequencing protocols—such as NanoSeqV1, CODEC-HpyCH4V, and HiDEF-seq—that rely on enzyme-based fragmentation and are limited to partial genome coverage, UDSeq leverages random fragmentation to achieve near-complete genome and exome representation. This enzyme-independent approach expands applicability across species and simplifies targeted sequencing without requiring protocol modifications. Additionally, our optimized library preparation pipeline enhances library conversion efficiency, generating up to four times more duplex molecules than NanoSeqV2 from the same DNA input. Combined with its compatibility with low-input samples and streamlined workflow, UDSeq offers both technical performance and cost-effectiveness, making it particularly valuable for studies involving scarce clinical material or environmental specimens.

In this study, we demonstrated the power of UDSeq across diverse applications. *In vitro*, it captured carcinogen-induced mutational signatures in heterogeneous cell populations—without the need for laborious clonal expansion. *In vivo*, it revealed exposure-specific mutation patterns in rodent models and enabled genome-wide mutation detection in non-model organisms including chickens and sheep. In human tissue biopsies, UDSeq recovered expected organ-specific mutational signatures and identified differences in mutational burden, further validating its utility for studying tissue-specific mutagenesis. While some of these applications could, in principle, be addressed with other duplex sequencing protocols, to the best of our knowledge UDSeq is the only method optimized to perform all of them, with experiments conducted across different laboratories confirming its versatility and with the protocol streamlined to facilitate adoption and use by others.

While UDSeq has demonstrated versatility across a wide range of applications and offers clear advantages over existing duplex sequencing protocols, it still shares certain limitations inherent to all short-read duplex sequencing. Specifically, the protocol is not well-suited for detecting large structural variants, complex rearrangements, or copy number alterations, which require long-range genomic context. Future integration with long-read sequencing technologies or complementary genomic platforms could overcome these challenges, further extending UDSeq’s utility to capture both small-scale mutations and large-scale genomic alterations.

In summary, UDSeq is a robust, scalable, and cost-efficient duplex sequencing protocol that enables accurate detection of rare somatic mutations at single-molecule resolution. Its flexibility across species, compatibility with limited input material, and high technical fidelity position UDSeq as a powerful tool for advancing studies of mutagenesis, somatic mosaicism, aging, cancer biology, and environmental exposures. Importantly, we provide a clear and streamlined protocol (**Supplementary Note 1**) that is easy to use and has been extensively validated across multiple independent laboratories, ensuring broad reproducibility and accessibility.

## METHODS

### Human biospecimens

All human biospecimens were collected with informed consent from all human research participants or their families. The tissue samples used in this study were collected post-mortem from deceased human participants by LIBD, not from living individuals. The collection was conducted in accordance with applicable national and state Institutional Review Board (IRB) regulations (study number: 1126332; IRB tracking number: 20111080). Sperm samples were collected from healthy ethnically diverse males enrolled according to approved human subjects’ protocols from the Institutional Review Board (IRB) of the University of California for blood, saliva, and semen sampling (140028, 161115). Genomic DNA was extracted using the DNeasy Blood and Tissue kit (QIAGEN, Cat# 69506, Valencia, CA) following the manufacturer’s recommendations.

### Cytotoxicity assessment

Cytotoxicity assessment was performed for all *in vitro* experiments. Specifically, cell viability was determined using the CellTiter-Glo® Luminescent Cell Viability Assay (Promega, Cat# G7572, Madison, WI), which quantifies ATP as an indicator of metabolically active cells. The reagent was added to each well of a 96-well plate at a 1:10 ratio. After a 10-minute incubation at room temperature, luminescence was recorded using a Cytation 5 Cell Imaging Multi-Mode Reader (BioTek, Winooski, VT). Relative cell viability was calculated as the percentage of luminescent signal from treated cells compared to untreated controls

### *In vitro* experiments

HepG2 human liver cancer cell line, derived from a well-differentiated hepatocellular carcinoma were purchased from ATCC (HB-8065). An hTERT immortalized non-cancerous human keratinocyte cell line (i.e., N/TERT-1) was purchased from Cellosaurus (RRID: CVCL_CW92). Normal oral keratinocytes (NOK) cell lines were a kind gift from Dr. Paul Lambert (University of Wisconsin-Madison, United States of America). The cells were generated by retroviral insertion of the human hTERT gene in oral epithelial cells derived from gingival tissue. The cells were propagated in the keratinocyte growth medium 2 (PromoCell GmbH, Heidelberg, Germany) and 1% penicillin/streptomycin. All other cells were cultured by following the recommended cell maintenance process from manufacturer using T25 (Thermo Fisher, 169900) or T75 (Thermo Fisher, 156800) flasks. Following cytotoxicity assessment, half-maximal inhibitory concentration (IC_50_) of environmental carcinogens was used for exposure with a specific duration of time. For *in vitro* experiments for profiling with UDSeq, no single cell clonal bottlenecking/passaging was done after exposure. For each experiment, cells were passaged only once after exposure, followed by DNA extraction. Following treatment, genomic DNA extraction was performed using the DNeasy Blood & Tissue Kit (QIAGEN, Cat# 69506, Valencia, CA), including RNase A treatment to eliminate RNA contamination. DNA concentrations were measured using the Qubit™ dsDNA Broad Range Assay Kit (Thermo Fisher Scientific, Cat# Q32850).

For *in vitro* experiment with bottlenecking, clonal expansion and profiling with bulk sequencing, primary human cells derived from human foreskin fibroblasts (HFFs) were passaged and clonally expanded by following the methods in Zhivagui *et al.*^56^. Cells were washed weekly, until clones reached confluency and were transferred progressively to T-75 flasks. 4NQO exposure (0.5 µM for 4 hours) was performed following cytotoxicity assessment to determine the IC_50_ concentration. Following exposure, cells underwent an additional clonal passage for ∼35 rounds of cell division, after which DNA was extracted and subjected to bulk whole-genome sequencing using the NEBNext® Ultra™ II DNA Library Prep Kit for Illumina® (E7645S). Clonal expansion results for other cells were based on previously generated sequencing data as reported in the original publications.

### *In vivo* experiments

Male SKH-1 mice (21–25 days old) were purchased from Charles River Laboratories (Wilmington, MA). These studies were performed under an approved Institutional Animal Care and Use Committee (IACUC) protocol 25-201636-HSC at the University of New Mexico. Mice were either controls (i.e., unexposed to ssUVR) or exposed to ssUVR (14 kJ/m^2^; ∼0.5 minimal erythema dose) 3 times per week for 30 weeks. Animals were sacrificed 4 weeks after the last ssUVR treatment. Animals were euthanized using CO_2_ followed by cervical dislocation and tissues were collected. Skin tissue was collected in 10% neutral buffered formalin, RNAlater, snap-frozen, and epidermal scrapings obtained from both ventral and dorsal skin. We have complied with all relevant ethical regulations for animal use. Genomic DNA extraction was performed using the DNeasy Blood & Tissue Kit (QIAGEN, Cat# 69506, Valencia, CA), including RNase A treatment to eliminate RNA contamination. DNA concentrations were measured using the Qubit™ dsDNA Broad Range Assay Kit (Thermo Fisher Scientific, Cat# Q32850).

F344 rats (21–25 days old) were purchased from Charles River Laboratories (Wilmington, MA). These studies were performed under an approved Institutional Animal Care and Use Committee (IACUC) protocol (#1802-35549A) at University of Minnesota. Following one week of acclimation, rats were treated with NNK (5 parts per million in drinking water) and were euthanized after 15 weeks. Control rats were provided with normal drinking water. Animals were euthanized using CO_2_ followed by cervical dislocation and tissues were collected. Lung tissues were from both control and NNK-exposed rats. Tissues were collected and flash frozen. Genomic DNA extraction was performed using the DNeasy Blood & Tissue Kit (QIAGEN, Cat# 69506, Valencia, CA), including RNase A treatment to eliminate RNA contamination. DNA concentrations were measured using the Qubit™ dsDNA Broad Range Assay Kit (Thermo Fisher Scientific, Cat# Q32850).

Chicken and sheep organs were obtained from a butcher shop in San Diego. Genomic DNA extraction was performed using the DNeasy Blood & Tissue Kit (QIAGEN, Cat# 69506, Valencia, CA), including RNase A treatment to eliminate RNA contamination. DNA concentrations were measured using the Qubit™ dsDNA Broad Range Assay Kit (Thermo Fisher Scientific, Cat# Q32850).

### UDSeq Library Preparation

The complete step-by-step UDSeq protocol is provided in **Supplementary Note 1**. Briefly, to minimize DNA damage during fragmentation, intact genomic DNA was enzymatically fragmented using NEBNext dsDNA Fragmentase (M0348S) or UltraShear (M7634L) to achieve an average fragment size of ∼350 bp. Fragmentation conditions were carefully optimized for each species and sample type. For human samples, both sperm and cell lines were fragmented for 15 minutes, while human tissue required 20 minutes. Mouse cell lines were also fragmented for 20 minutes, but mouse tissue needed a longer duration of 25 minutes. Similarly, rat tissues were fragmented for 25 minutes to achieve optimal results. Fragmented DNA was then used for UMI adapter ligation with the xGen™ cfDNA & FFPE DNA Library Preparation Kit. All steps were carried out on magnetic beads to reduce DNA loss during purification, thereby improving library conversion efficiency (**Supplementary Figure 1b**). In the final step, an appropriate femtomole input amount was used for PCR amplification to incorporate sample index sequences compatible with Illumina® sequencing platforms.

### DNA Quantification, Dilution, and PCR Amplification

A key strength of UDSeq lies in the accurate quantification of adapter-ligated DNA using qPCR. To avoid the variability introduced by mixed primer sets during quantification, we utilized NEBNext® Library Quant DNA Standards, which reliably quantify UMI-ligated molecules. For size correction, we used 330 bp for the standards. For adapter-ligated DNA, we estimated fragment size by adding 82 bp (accounting for UMI-containing adapters) to the average fragment length determined by TapeStation. For example, a sample with an average fragment size of 370 bp was quantified using 452 bp as the effective fragment length. Additional details are provided in **Supplementary Note 1**.

For library amplification, we used 0.2 fmol of input DNA and 15 PCR cycles to achieve ∼90× whole-genome coverage in human samples. For mouse samples, we used 0.15 fmol with the same number of cycles. For other species, input amounts were adjusted as appropriate to target ∼80% duplicated and ∼20% unique reads (**Supplementary Note 1)**.

Targeted hybrid capture was performed using 6–8 multiplexed samples per reaction, with 500 ng of adapter-ligated DNA per capture. The complete targeted capture protocol, including exome and panel-based enrichment, is described in the **Supplementary Note 1**. The pre-made UDSeq libraries were sequenced on an Illumina NovaSeq 6000 and NovaSeq X platform using 150 PE sequencing chemistry to effective data volume.

### Trimming, Alignment, and Mutation Identification

All bioinformatics analyses were performed within the Triton Shared Compute Cluster *(San Diego Supercomputer Center (2022): Triton Shared Computing Cluster. University of California, San Diego. Service.* https://doi.org/10.57873/T34W2R). Somatic mutations and mutational burden from UDSeq data with matched normal were analyzed using DupCaller^57^ ver1.0.1. Briefly, Paired-end FASTQ files with equal-length barcodes at the start of each read were preprocessed to remove barcodes and align sequences to the reference genome using BWA. PCR and optical duplicates were marked using GATK. DupCaller constructs sample-specific error profiles by analyzing single-strand mismatches and single-read discrepancies. These profiles are stratified by trinucleotide context and homopolymer length for substitutions and indels, respectively. A strand-aware probabilistic model calculates genotype likelihoods and assigns confidence scores to candidate mutations. Mutations exceeding a confidence threshold are retained. Post-calling filters were used to exclude low-quality reads, common germline variants, and noisy loci.

### Mutational profile and signature analysis

The variant call format files (VCFs) from DupCaller were used for mutational profiles and signatures assignment. Analysis of mutational profiles was performed using our previously established methodology with the SigProfiler suite of tools. Briefly, mutational matrices for SBS, DBS and Indels were generated with SigProfilerMatrixGenerator^58^ (Version 1.2.16). Plotting of each mutational profile was done with SigProfilerPlotting (Version 1.3.13). Assignment of mutational signatures to samples was done with SigProfilerAssignment^59^. Mutational profile of sperm samples from NanoSeqV1^35^ and HiDEF-Seq^38^ were obtained from their corresponding publications. Parental *de novo* mutations were obtained from *Halldorsson, B. V. et al.*,^41^. and the patterns of the mutations are plotted with SigProfilerMatrixGenerator. Regression plots mutation rate was calculated as previously described in Ref. ^38^. The corrected mutation burdens output from DupCaller was used for plotting using R statistical language^60^.

## Data availability

All whole-genome sequencing data have been or will be deposited in the Sequence Read Archive (SRA) or the database of Genotypes and Phenotypes (dbGaP), as appropriate. Duplex sequencing data from N/TERT-1 and HepG2 cell lines, as well as SKH-1 mouse, F344 rat, sheep, and chicken tissues, are available under accession number PRJNA1262723. Duplex sequencing data for NOK cells are accessible via PRJNA1196807. Bulk clonal expansion sequencing data for iPSC, BEAS-2B, and N/TERT-1 were obtained from the respective publications cited in the manuscript. Data for human foreskin fibroblasts (HFF), generated as part of this study, are also deposited under PRJNA1262723. Whole-genome sequencing data from human subjects will be made available in dbGaP upon acceptance of the manuscript. Patient ID 7614 data can be accessed via PRJNA799597. Sequencing data for sperm samples are available under accession numbers PRJNA660493, PRJNA753973, and PRJNA588332. All other data are available from the corresponding authors or other sources upon reasonable request.

## ACKNOWLEDGMENTS

The authors would like to thank Cécilia Sirand for her technical support in performing some of the cell line experiments and Dr Fekadu Kassie for assistance with the NNK rat study. This work was supported by the US National Institute of Health grants R01ES032547, R01ES036931, R01CA269919, R01CA296974, P01CA281819, and U01CA290479 to L.B.A. and RO1CA220376 to S.B. as well as by L.B.A.’s Packard Fellowship for Science and Engineering and the UC San Diego Sanford Stem Cell Institute. The work presented here is also supported by a network grant from The Larry L. Hillblom Foundation to L.B.A. and J.G.G. as well as by UK Grand Challenge 2016 Award “Mutographs of Cancer” C98/A24032 to L.B.A. and J.Z. This work was supported in part by NIH award R00HD111686 to X.Y. The computational analyses reported in this manuscript have utilized the Triton Shared Computing Cluster at the San Diego Supercomputer Center of UC San Diego. The funders had no roles in study design, data collection and analysis, decision to publish, or preparation of the manuscript.

## DISCLAIMER

Where members are identified as personnel of the International Agency for Research on Cancer/World Health Organization, the authors alone are responsible for the views expressed in this article and they do not necessarily represent the decisions, policy or views of the International Agency for Research on Cancer/World Health Organization.

## AUTHOR CONTRIBUTIONS

S.P.N. and L.B.A. conceptualized the study and designed the UDSeq protocol with assistance from Y.C and advice from M.P., J.Z., and J.G.G. S.P.N. optimized the protocol and performed comparison with assistance and advice from S.A., S.S., A.K., N.S., I.R.S., Z.J., L.C., and M.Z. Access and analysis of human samples was performed by S.P.N., Y.C., X.Y., and J.G.G. Access and analysis of mouse samples was performed by S.P.N., Y.C., R.M.S., L.G.H., and K.J.L. Access and analysis of rat samples was performed by S.P.N., Y.C., F.C.J, and S.B. Access and analysis of cell line experiments was performed by S.P.N., Y.C., B.C., M.K., and J.Z. S.P.N. and L.B.A. wrote the manuscript with input from all co-authors. All authors read and approved the final manuscript.

## SUPPLEMENTARY FIGURE LEGENDS

**Supplementary Figure 1: UDSeq protocol overview and comparative performance.**

***(a)*** Detailed schematic of the UDSeq protocol, comprising four major steps: *(i)* DNA extraction and fragmentation, *(ii)* end repair and adapter ligation for library preparation, *(iii)* library quantification and sequencing, and *(iv)* data analysis. ***(b)*** Comparative overview of genomic coverage achieved for human cortex (UDSeq), human cortex (NanoSeqV1), and human blood (bulk WGS) samples across different target regions (exome and genome) at varying coverage thresholds. ***(c)*** Quantitative polymerase chain reaction (quantitative PCR) amplification curves and quantification of unique molecular identifier–ligated molecules (in femtomoles) demonstrating library conversion efficiency of Universal Duplex Sequencing (UDSeq) versus NanoSeqV2 using 10 nanograms of fragmented DNA input. UDSeq achieved approximately four-fold higher conversion efficiency. Horizontal lines indicate medians; boxes represent interquartile ranges (IQR), and whiskers extend to 1.5× IQR. Statistical significance was assessed using a two-sided t-test (p=0.00022). ***(d)*** Cost-effectiveness comparison of UDSeq, NanoSeq, CODEC, and HiDEF-seq. Projected error rates and estimated cost per megabase of duplex coverage for CODEC (using HpyCH4V enzymatic fragmentation), CODEC (sonication), NanoSeqV1, HiDEF-seq, and UDSeq. Error rates are shown on the y-axis.

**Supplemental Figure 2: Mutational profiles and genomic analyses across species and tissues. *(a)*** SBS-96 mutational profiles from distinct anatomical layers of the kidney in three individual sheep, with the relative contributions of the clock-like COSMIC signatures SBS1 and SBS5 shown alongside each profile. Each SBS-96 profile represents all single-base substitutions (C>A, C>G, C>T, T>A, T>C, T>G) within their trinucleotide sequence context. Y-axis scales are adjusted independently to optimize visualization of mutation percentages within each trinucleotide context. ***(b)*** SBS-96 mutational profiles from skin, breast, and pancreas tissues in chicken, with the contributions of the clock-like COSMIC signatures SBS1 and SBS5, as well as the reactive oxygen species–associated signature SBS18, displayed adjacent to each profile. Profiles are displayed in a format consistent with *(a)*. ***(c)*** Box plots showing mutation burden per base pair (log scale) for kidney cortex and two replicates of kidney medulla from sheep, as well as for skin, breast, and two replicates of pancreas from chicken. Horizontal lines indicate medians; boxes represent interquartile ranges (IQR), and whiskers extend to 1.5× IQR. ***(d)*** Bar plots showing the percentage of whole-genome duplex coverage achieved for chicken, sheep, rat, and mouse using the UDSeq approach. Coverage is expressed as the proportion of the genome successfully sequenced at the desired depth.

