## Supplementary figures and images for "A Universal Duplex Sequencing Approach for Accurate Detection of Somatic Mutations"

### Figure S1

#### Supplementary figure 1

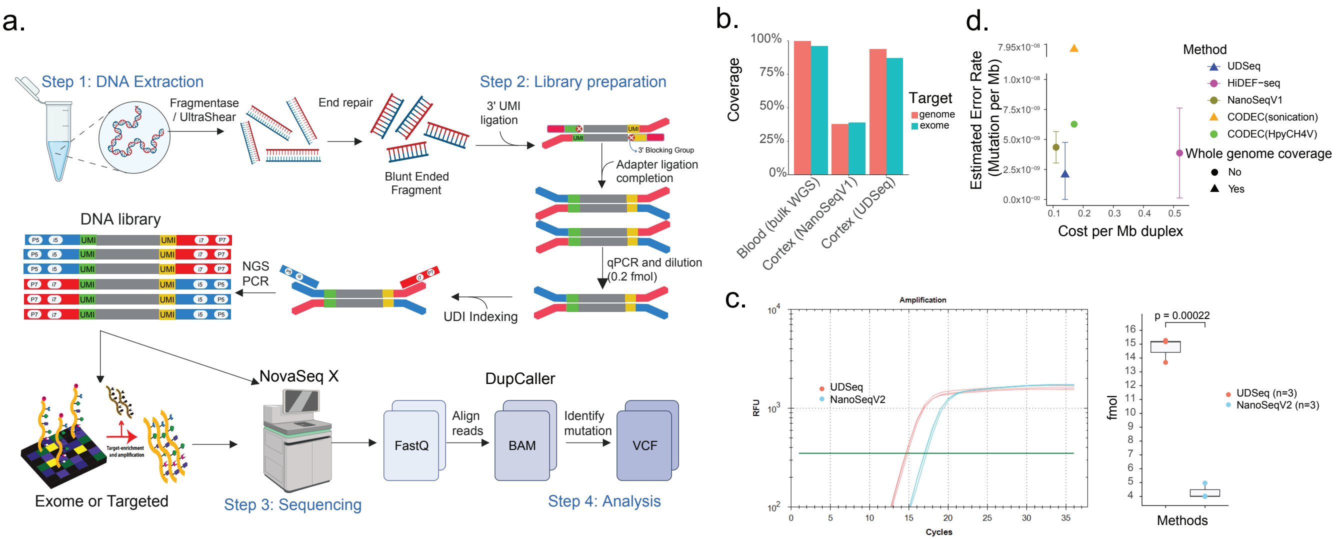

### Figure S2

Supplementary figure 2

a.

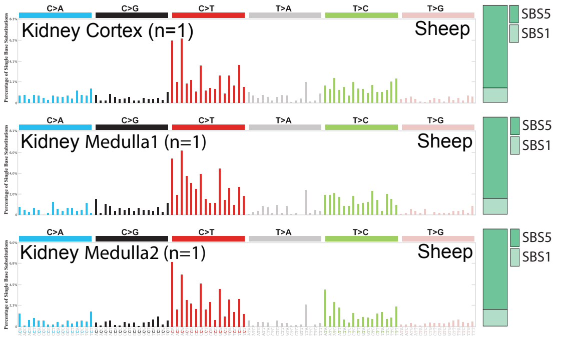


c.

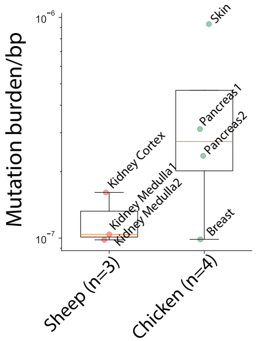


b.

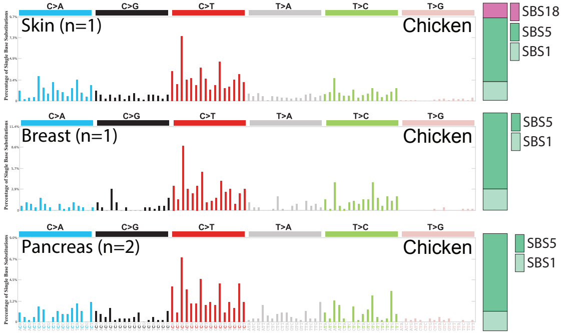


d.

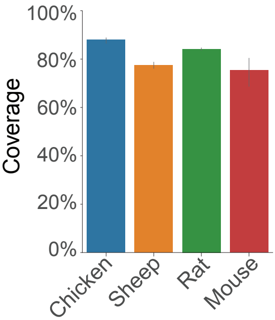
