## Supplementary Note 1 for "A Universal Duplex Sequencing Approach for Accurate Detection of Somatic Mutations"

#### Version History

| Version | Release Date | Description |
|---|---|---|
| 1.0.0 | September 5, 2025 | First edition |

### TABLE OF CONTENTS


Required Materials ii

**UDSeq WGS Protocol**

- A. Library Preparation 1
- B. DNA Quantification by qPCR 3
- C. Indexing and NGS-PCR 4

**Exome/Targeted UDSeq: Tube Protocol**

- A. Hybridization Reaction 5
- B-F. Capture & Washes 6
- G. Post-Capture PCR & Final Steps 8

Appendix I – Bead Purification Protocols 9

Appendix II – WGS Regular Coverage Calculations 10

Appendix III – Hybridization Input Requirements for Targeted Duplex Coverage 10

Appendix IV – Examples of Expected Results 10


#### Key to Symbols

**Pre NGS-PCR Area**
Amplicon-free environment.

**Reaction on Beads**
Indicates a step performed on beads.

**Safe Stopping Point**
Protocol can be safely paused.

**Tip**
Helpful advice for success.

### Required Materials

#### Reagents for UDSeq WGS

- NEBNext® dsDNA Fragmentase® (M0348L) or NEBNext UltraShear® (M7634L)
- xGen™ cfDNA & FFPE DNA Library Preparation Kit (10010203 or 10010207)
- Nuclease-free water (NFW) (Integrated DNA Technologies or IDT; 11-05-01-14)
- 1X IDTE buffer pH 8.0 (IDT; 11-05-01-13)
- 0.5 M EDTA (Thermo Fisher, 15575020)
- Agencourt AMPure XP beads (Beckman Coulter: A63882)
- 2.5M Sodium Chloride, 20% PEG Solution. 500mL, Sterile. (NaCl+PEG) (Teknova, P4137)
- 75% Ethanol (freshly made from 100% Ethanol)
- iTaq Universal SYBR Green Supermix (2x) (1725124)
- NEBNext® Library Quant DNA Standards (E7642S)
- qPCR primers:

  Forward primer: 5'-ACACTCTTTCCCTACACGAC-3'
  Reverse primer: 5'-GTGACTGGAGTTCAGACGTG-3'
- NEBNext® Ultra™ II Q5® 2X Master Mix (NEB; M0544X)
- UDI containing PCR primers:

  i5: 5'-AATGATACGGCGACCACCGAGATCTACAC[barcode]ACACTCTTTCCCTACACGACGCTCTTCCGATC*T-3'
  i7: 5'-CAAGCAGAAGACGGCATACGAGAT[barcode]GTGACTGGAGTTCAGACGTGTGCTCTTCCGATC*T-3'

  *phosphorothioate bond
- Qubit 1X dsDNA HS Assay Kit (Thermo Scientific, Q33231)
- Agilent High Sensitivity DNA Kit (to be used with a TapeStation or Bioanalyzer instrument)

#### Additional Reagents for UDSeq Targeted Protocol

- xGen™ Exome Hyb Panel v2 (10005153) or xGen™ Pan-Cancer Hybridization Panel (1056204)
- xGen™ Hybridization and Wash Reagents v2 (10010351)
- xGen™ Hybridization and Wash Beads v2 (10010353)
- xGen™ Universal Blockers TS, 16 rxn (1075474)
- xGen™ Library Amplification Primer Mix (1077675)
- xGen™ Human Cot DNA, 150 µL (1080768)
- Full length library amplification/ quantification primers:

  P5: 5'-AATGATACGGCGACCACCGA-3'
  P7: 5'-CAAGCAGAAGACGGCATACGA-3'

#### Equipments

- Full set of pipettes and tips
- Hard-Shell Low-Profile Skirted and Semi-Skirted PCR Plates
- Thermal cycler (e.g., Bio-Rad T100).
- Qubit™ 4 Fluorometer (Invitrogen)
- Plate magnet/12 well magnet
- Real-Time PCR Detection System (CFX Connect, Bio-Rad)
- TapeStation System (Agilent, 4150 or 4200)
- NovaSeq 6000 or NovaSeq X Sequencing System

### UDSeq WGS Protocol

**All steps should be performed in a clean, pre-PCR work area.**

#### A. Library Preparation

1. **DNA Clean-up:** Equilibrate AMPure XP beads to room temperature. Per sample, mix **50 µL** AMPure XP beads with **50 µL** NFW. Add **100 µL** of this 50:50 mix to each 20 µL (~10-200 ng) sample. Mix well by pipetting up and down and allow DNA to bind to beads, wash 2x with 75% EtOH (see Appendix I below) and resuspend the beads in either **16 µL** or **26 µL** 1X IDTE buffer depending on fragmentation of choice.

**TIP:** For beads purification, see details at the end. Use freshly (same day) prepared 75% ethanol for washing the beads for each purification.

**TIP:** If DNA concentration is low, adjust volumes as needed: 50 µL AMPure XP beads, xx µL DNA (to reach max ng), and xx µL 1X TE (to bring total to 120 µL)

2. **Fragmentation:**

Choose one of the following fragmentation methods:

<table>
<tr><th>Fragmentase</th><th>UltraShear</th></tr>
<tr>
<td>
1x Mix (20 µL Total):
<ul>
<li>16.00 µL DNA</li>
<li>2.00 µL Reaction Buffer V2</li>
<li>2.00 µL Fragmentase Enzyme</li>
</ul>
<table>
<tr><th colspan="2">Thermocycler Program (Lid heat off)</th></tr>
<tr><td>37°C</td><td>20 minutes</td></tr>
<tr><td>4°C</td><td>Hold (∞)</td></tr>
</table>
Add 5 µL of 0.5M EDTA (inactivation), mix and add 25 µL of NFW (Final volume: 50 µL).
</td>
<td>
1x Mix (44 µL Total):
<ul>
<li>26.00 µL DNA</li>
<li>14.00 µL Reaction Buffer</li>
<li>4.00 µL UltraShear Enzyme</li>
</ul>
<table>
<tr><th colspan="2">Thermocycler Program (Lid @ 75°C)</th></tr>
<tr><td>37°C</td><td>20 minutes</td></tr>
<tr><td>65°C</td><td>15 minutes (inactivation)</td></tr>
<tr><td>4°C</td><td>Hold (∞)</td></tr>
</table>
</td>
</tr>
</table>

3. **Clean-up:** Perform a 1.8x purification by adding **50 µL** NaCl+PEG solution and **40 µL** AMPure XP beads to your 50 µL sample. Resuspend in **50 µL** 1X IDTE buffer.

**TIP:** Check the fragment size on TapeStation. Acceptance criteria is 350-500 bp peak with <10% at >600 bp. If >10% fragments >600 bp, perform 0.65–0.7x right-side cleanup. A small fragment peak <350 bp will result in less uniform coverage of the genome. Optimization required for different species of interest and different tissue types.

**Safe Stopping Point (Optional): Store fragmented DNA at 4°C overnight or -20°C for long term.**

4. **End Repair** (IDT, 10010203 or 10010207):

Add **9 µL** of End Repair Mix to each 50 µL of fragmented DNA. Final volume is 59 µL. Incubate at 20˚C for 30 minutes on a thermocycler (lid heat off).

| End Repair Mix (9 µL Total) | |
|---|---|
| End Repair Buffer | 6.00 µL |
| End Repair Enzymes | 3.00 µL |

5. **Ligation1** (IDT, 10010203 or 10010207):

Prepare Ligation1 Mix. This ligation mix is going to be used as resuspension solution after beads purification.

| Ligation1 Mix (30 µL Total) | |
|---|---|
| Ligation1 Buffer | 25.00 µL |
| Ligation1 Adapter | 2.00 µL |
| Ligation1 Enzyme | 3.00 µL |

Add **100 µL** NaCl+PEG to each 59 µL sample. Mix well by pipetting up and down and allow DNA to bind to beads (10 minutes) and wash 2x with 75% EtOH (see Appendix I below). Resuspend beads in **30 µL** of Ligation1 Mix.

Perform Ligation1 (on beads) by incubating with the following program (lid heat @ 70°C):

| Step | Temperature | Time |
|---|---|---|
| 1 | 20°C | 15 minutes |
| 2 | 65°C | 15 minutes |
| 3 | 4°C | Hold (∞) |

6. **Ligation2** (IDT, 10010203 or 10010207):

Prepare Ligation2 Mix:

| Ligation2 Mix (10 µL Total) | |
|---|---|
| Ligation2 Buffer | 4.50 µL |
| Ligation2 Adapters | 4.00 µL |
| Ligation2 Enzyme A | 0.50 µL |
| Ligation2 Enzyme B | 1.00 µL |

Add **10 µL** of Ligation2 Mix to the 30 µL reaction from Ligation1 and mix with pipetting. Incubate at 65°C for 30 min on a thermocycler (lid heat @ 70°C).

7. **Post-ligation Clean-up:** Add **100 µL** NaCl+PEG to each 40 µL sample beads mixture. Mix well by pipetting up and down and allow DNA to bind to beads, wash 2x with 75% EtOH and elute in **50 µL** 1X IDTE buffer.

**Safe Stopping Point (Optional): Store fragmented DNA at 4°C overnight or -20°C for long term.**

### B. DNA Quantification by qPCR

8. **Prepare for qPCR:** Quantify adapter-ligated DNA using qPCR with the NEB Library Quantification Standard (NEB#E7642S) to determine the optimal number of amplification cycles prior to Illumina cluster generation.

Sample Dilution: Dilute each sample 1:500 and/or 1:1000 in 1X IDTE buffer.

qPCR Setup: Prepare triplicate 10 µL reactions in a 96- or 384-well plate: 8 µL of qPCR master mix 2 µL of diluted sample or standard.

Reagents: Use any suitable library quantification master mix. For Bio-Rad qPCR systems, iTaq™ Universal SYBR® Green Supermix has shown optimal performance.

| 1x mix (8 µL) | |
|---|---|
| iTaq Universal SYBR Green Supermix (2x) | 5.00 µL |
| qPCR_F primer (10 µM) | 0.20 µL |
| qPCR_R primer (10 µM) | 0.20 µL |
| NFW | 2.60 µL |

9. **qPCR Reaction:** Run on qPCR system using a program with a 3-step cycle and melt curve analysis.

qPCR plate setup example:

| | 1 | 2 | 3 | 4 | 5 | 6 |
|---|---|---|---|---|---|---|
| **A** | NTC* | NTC | NTC | Sample 1 | Sample 1 | Sample 1 |
| **B** | Std-1 | Std-1 | Std-1 | Sample 2 | Sample 2 | Sample 2 |
| **C** | Std-2 | Std-2 | Std-2 | Sample 3 | Sample 3 | Sample 3 |
| **D** | Std-3 | Std-3 | Std-3 | Sample 4 | Sample 4 | Sample 4 |
| **E** | Std-4 | Std-4 | Std-4 | Sample 5 | Sample 5 | Sample 5 |
| **F** | Std-5 | Std-5 | Std-5 | Sample 6 | Sample 6 | Sample 6 |
| **G** | Std-6 | Std-6 | Std-6 | Sample 7 | Sample 7 | Sample 7 |
| **H** | IC* | IC | IC | Sample 8 | Sample 8 | Sample 8 |

NTC* = No Template Control; IC* = Internal Control (optional)

qPCR reaction setup. Run samples on a qPCR machine e.g., Bio-Rad CFX connect qPCR system.

| Step | Temperature | Time | Cycles |
|---|---|---|---|
| 1 | 95°C | 3 minutes | 1 |
| 2 | 95°C | 30 seconds | **35 Cycles** |
| 3 | 60°C | 30 seconds[1] | |
| 4 | 65-95°C | Melt curve analysis | - |
| 5 | 4°C | ∞ | 1 |

[1] Increase to 60 seconds for long-insert libraries (>700 bp).

10. **Quantification Analysis:**

Calculate nM concentration using the standard curve from the Bio-Rad PCR analysis software and the library size from the Agilent TapeStation.

**TIP:** qPCR acceptance criteria: standard curve $R^2$ ≥0.980, uniform peaks in the melt curve, triplicate Cq SD ≤ 0.3, and NTC is a flat line (minimal amplification). Quantification efficiency ≥95%.

### C. Indexing and NGS-PCR

11. **Perform quantification analysis:** You can use kappa library quantification template according to the library dilution. Determine the nM (fmol/ µL) concentration per adapter-ligated library as follows:.

$$[nM] = \frac{\text{qPCR conc. (pM)}}{1000} \times \frac{\text{Size of qPCR standard (330 bp)}}{\text{Avg. fragment size}} \times \text{dilution factor}$$

Example: If qPCR reports the sample's average concentration is 0.5 pM for its 1:500 dilution, and the sample's average fragment size from TapeStation is 400 bp, the calculation is:

$$\frac{0.5}{1000} \times \frac{330}{400} \times 500 = 0.20625 \text{ nM}$$

12. **Dilute your adapter-ligated library** to your desired fmol input amount in **20 µL** using 1X IDTE buffer to achieve duplicate rates close to the optimal 80%. The fmol amount needs to be optimized to achieve 80% duplication rate. Refer to step 14 and Appendix II for guidance.

    **Sequencing of matched normal (if needed)** Matched normal is an optional parameter for targeted UDSeq, but it is recommended for WGS UDSeq. If bulk sequenced matched normal samples from the same patients or experiment are not obtainable, matched normal samples can still be sequenced at lower cost by using undiluted UDSeq WGS libraries. ≥2 fmol and 11 total cycles of NGS-PCR of library sequenced at 30X genome equivalent is enough to provide ~30X matched normal genome-wide coverage.

13. **Indexing with NGS-PCR**: First, prepare a 25 µL sample mix containing **20 µL** of your diluted DNA and 5 µL of the barcoded UDI primers (**2.5 µL** of i5 and **2.5 µL** of i7). Then, add **25 µL** of the NEBNext Ultra II Q5 2X master mix to this sample mix for a final volume of 50 µL.

    Preparation of PCR mix:

| 1x indexing mix (50 µL) | |
|---|---|
| Adapter-ligated DNA (diluted in 1X IDTE buffer) | 20.00 µL |
| i5 primer (UDI, 10 µM) | 2.50 µL |
| i7 primer (UDI, 10 µM) | 2.50 µL |
| NEBNext Ultra II Q5 2X master mix | 25.00 µL |

**TIP:** When handling barcoded UDI primers, be especially careful not to cross-contaminate. Change gloves frequently and open one tube at a time.

14. **NGS-PCR set up:** Place the sample mix into a thermocycler with lid heated to 105°C.

| Steps | Temp | Time | Cycles |
|---|---|---|---|
| 1 | 98°C | 30 seconds | 1 |
| 2 | 98°C | 10 seconds | 10-15<br>Refer to Appendix II |
| 3 | 65°C | 75 seconds | |
| 4 | 65°C | 5 minutes | 1 |
| 5 | 4°C | ∞ | - |

15. **Purify twice with magnetic beads** using 0.65x sample volume of AMPure XP beads and elute in 50 µL 1X IDTE buffer.
16. **Quantify each sample** with Qubit high sensitivity dsDNA quantification kit (Invitrogen).
17. **Pool samples** together for sequencing.

18. **Run pool on BioAnalyzer/TapeStation** and adjust the pool concentration to check the final library.

$$[nM] = \frac{\text{Conc. of DNA from Qubit } (\frac{ng}{\mu L})}{660 \times \text{Avg. fragment size (bp)}} \times 1 \cdot 10^6$$

19. **Sequence on preferred Illumina sequencer**. Use unique dual indices. It is recommended to add 1–5% PhiX if you don't have a complex enough pooled library (if you pooled less than 6 samples together). Follow platform-recommended cluster densities for NovaSeq X Plus 25B using 150 PE read lengths, or the utilized sequencing platform.

### Exome/Targeted UDSeq: Tube Protocol

Note: Starting DNA concentration for targeted capture is **≥50 ng** of fragmented DNA. If doing exome capture, take **≥10 fmol** of UMI ligated DNA from step 11 and perform 15 cycle NGS-PCR from step 14 in the previous page. If doing targeted capture, take **≥30 fmol** of UMI ligated DNA from step 11 and perform 12 cycle NGS-PCR from step 14 in the previous page. Refer to Appendix III for more details.

1. Before you start, create the two cycling programs:

**Hybridization program (HYB, lid at 100°C)**

| Steps | Temperature | Time | Volume |
|---|---|---|---|
| 1 | 95°C | 30 seconds | 17 µL |
| 2 | 65°C | 4 hours | |
| 3 | 65°C | ∞ | |

**WASH program (lid at 70°C)**

| Steps | Temperature | Time | Volume |
|---|---|---|---|
| 1 | 65°C | ∞ | 34 µL |

2. Thaw xGen Hyb Panels at room temperature.

#### A. Hybridization Reaction (if pooling, 6 samples) (~5 hours).

**TIP:** Use of SpeedVac is recommended for dry down process to increase the final yield.

3. Add **500 ng** of one adapter-ligated library to a 1.5 mL tube. If multiplexing, pool 500 ng of each library into the sample tube.
4. Add **7.5 µL** Human Cot DNA and 1.8x AMPure XP beads.
5. Briefly vortex and incubate for 10 minutes at room temperature. See step 7 on how to prepare the Hybridization Reaction Mix.
6. Place tube on magnet until supernatant is clear (~2 minutes) and discard the supernatant. Wash 2x with 75% ethanol. Allow the beads to air dry for ~2 minutes until the beads are matte to remove any residual ethanol. Avoiding over-drying the beads.
7. Resuspend in **19 µL** of Hybridization Reaction Mix. Incubate for 5 minutes at room temperature, place on magnet for 5-10 minutes, and transfer **17 µL** of supernatant in another tube (avoid any beads carry over).

| 1x Hybridization reaction mix (19 µL) | |
|---|---|
| xGen 2X Hybridization Buffer | 9.5 µL |
| xGen Hybridization Buffer Enhancer | 3.0 µL |
| xGen Universal Blockers based on library adapters | 2.0 µL |
| xGen Hybridization Capture Panel V2 (or custom) | 4.5 µL |

8. Place the 17 µL sample tube in the thermal cycler and start the HYB program (will come back to this during bead capture). The final HYB volume of this singleplex or multiplex capture reaction is 17 µL.

#### B. Prepare Wash Buffers (start this process approximately 3 hours into the 4-hour HYB program)

9. Take out the Dynabeads™ M-270 Streptavidin from 4°C and leave at room temperature for ~30 minutes before wash.
10. Dilute the following buffers to create 1x working solutions:

| Buffers | NFW | Buffer | Total | Storage |
|---|---|---|---|---|
| xGen 2X Bead Wash Buffer | 160 µL | 160 µL | 320 µL | Room temperature |
| xGen 10X Wash Buffer 1 | 252 µL | 28 µL | 280 µL | Aliquot 110 µL into separate tube and heat to 65°C. Remaining solution at room temperature |
| xGen 10X Wash Buffer 2 | 144 µL | 16 µL | 160 µL | Room temperature |
| xGen 10X Wash Buffer 3 | 144 µL | 16 µL | 160 µL | Room temperature |
| xGen 10X Stringent Wash Buffer | 288 µL | 32 µL | 320 µL | Aliquot into two tubes (160 µL each). Heat tubes to 65°C |

If the 10X Wash Buffer 1 is cloudy, heat the bottle in 65°C water bath.

11. Prepare the following bead resuspension mix in a low-bind tube:

| 1x Bead resuspension mix (17 µL) | |
|---|---|
| xGen 2X Hybridization Buffer | 8.5 µL |
| xGen Hybridization Buffer Enhancer | 2.7 µL |
| NFW | 5.8 µL |

#### C. Prepare Dynabeads™ M-270 Streptavidin Beads for Capture

**TIP:** Only perform bead washes with beads that have equilibrated at room temperature.

12. Mix the Streptavidin beads thoroughly by vortexing for 15 seconds.
13. Aliquot **50 µL** per capture of Streptavidin beads into a single 0.2 mL low-bind tube (e.g., 1 capture= 50µL beads, 2 captures = 100µL beads).
14. Add **100 µL** per capture of 1X Bead Wash Buffer. Gently pipette to mix 10 times.
15. Place the tube on magnetic rack for ~1 minute. Remove and discard supernatant, keeping the beads in tube.
16. Remove the tube from the magnet.
17. Perform steps 14-16 twice for a total of **3 washes**.
18. Resuspend the beads in **17 µL** per capture of bead resuspension mix from step 11 and mix thoroughly. (Aliquot 17 µL of resuspended beads into a new low-bind 0.2 mL tube for each capture reaction if doing multiple captures).

#### D. Perform Exome Capture: Mix reaction from step A with step C

19. Place the 1X Wash Buffer 1 (110 µL aliquot) and the 1X Stringent Wash Buffer (both aliquots) in a 65°C water bath for at least 15 minutes.

**TIP:** Do this now so that they will be at the correct temperature during the heated washes.

20. After the 4-hr HYB incubation, take the tube out of the thermal cycler and stop the HYB program.
21. Immediately after the HYB program, start the WASH program.
22. Transfer **17 µL** of resuspended Streptavidin beads to the 0.2 mL tube containing the sample (HYB volume = 17 µL, final volume = 34 µL).
23. Vortex to ensure that sample is fully resuspended and briefly centrifuge (25 seconds).

24. Place sample tube in the thermal cycler and set a timer for 45 minutes.
25. Every 10-12 minutes, remove the tube from the thermal cycler and gently vortex to ensure the sample is fully resuspended.
26. At the end of the 45 minutes, take the sample off from the thermal cycler and immediately proceed to heated washes.

#### E. Heated Washes

27. Transfer **100 µL** of heated Wash Buffer 1 to the sample, then pipette mix 10 times, minimizing bubbles.
28. Place tube on magnetic rack for 1 minutes and remove supernatant.
29. Remove tube from magnet and add **150 µL** of heated Stringent Wash Buffer to the sample.
30. Pipette mix 10 times, minimizing bubbles.
31. Incubate in the water bath at 65°C for 5 minutes.
32. Place tube on magnet for 1 minute. Remove and discard supernatant and repeat step 29 to 32.

#### F. Room Temperature Washes

**TIP:** Vigorously mix the samples during the room temperature washes to ensure the beads are remain fully resuspended.

33. Remove and discard the supernatant. Add **150 µL** of Wash Buffer 1 equilibrated to room temperature.
34. Vortex thoroughly until fully resuspended and incubate for 2 minutes while alternating between vortexing for 30 seconds and resting for 30 seconds to ensure the mixture remains homogenous.
35. At the end of the incubation, briefly centrifuge the tube and place on magnet for 1 minute.
36. Remove and discard the supernatant.
37. Add **150 µL** of Wash Buffer 2 and repeat steps 34-36.
38. Add **150 µL** of Wash Buffer 3 and repeat steps 34-36.
39. With the sample tube still on the magnet, use a fresh pipette tip to remove residual Wash Buffer 3 from the tube, then remove the tube from the magnet.
40. Add **20 µL** of NFW to each capture and pipette mix 10x to resuspend any beads stuck to the side of the tube. *Do not discard beads.

#### G. Perform Post-Capture PCR

41. Choose one of the following master mixes and add the following components to create the Amplification Reaction Mix:

| NEBNext Ultra II | | IDT xGen | |
|---|---|---|---|
| **1x indexing mix (50 µL)** | | **1x indexing mix (50 µL)** | |
| Adapter-ligated DNA (diluted in 1X IDTE buffer) | 20.00 µL | Adapter-ligated DNA (diluted in 1X IDTE buffer) | 20.00 µL |
| P5 primer (10 µM) | 2.50 µL | xGen Library Amplification Primer Mix (20 µM) | 5.00 µL |
| P7 primer (10 µM) | 2.50 µL | xGen 2X HiFi PCR Mix | 25.00 µL |
| NEBNext Ultra II Q5 2X master mix | 25.00 µL | | |

42. Place the tube in a thermal cycler, and run the following program with the heated lid set at 105°C:

| Steps | Temp | Time | Cycles |
|---|---|---|---|
| 1 | 98°C | 45 seconds | 1 |
| 2 | 98°C | 15 seconds | 15 |
| 3 | 60°C | 30 seconds | |
| 4 | 72°C | 30 seconds | |
| 5 | 72°C | 1 minute | 1 |
| 6 | 4°C | ∞ | - |

**Safe Stopping Point (Optional): Store amplified captures at 4°C overnight or -20°C for long term.**

#### Post-capture PCR clean up

**TIP:** Ensure AMPure XP beads have been equilibrated to room temperature before proceeding.

43. Prepare fresh 75% ethanol
44. Add **75 µL** (1.5x) of AMPure XP beads to each amplified capture (transfer to larger tube if needed). After adding the beads, mix thoroughly and incubate for 5-10 minutes.
45. Place the sample tube on a magnet until the supernatant is clear (2-5 minutes). Remove supernatant without disturbing the beads.
46. While keeping the tube on the magnet, add **125 µL** of prepared 75% ethanol and ethanol wash twice.
47. Remove the sample tube from the magnet and elute in **22 µL** 1X IDTE buffer. Incubate 5 minutes. Place back onto the magnet and wait 1-2 minutes until supernatent is clear. Transfer **20 µL** to a fresh tube, discard beads.
48. Qubit and TapeStation to verify your yield.
49. Sequence at the desirable duplex depth. Refer to Appendix III.

**TIP:** For targeted capture, follow the same as exome capture, but replace the panel with your panel of interest. For our target of interest, we used xGen™ Pan-Cancer Hybridization Panel (1056205).

**TIP:** Matched normals are not needed for exome or targeted UDSeq. But you can still use matched normal during analysis if available.

### Appendix I: Bead Purification Protocols

**TIP:** Ensure AMPure XP beads have been equilibrated to room temperature before proceeding.

Two related but distinct purification procedures are used throughout the protocol. Please pay close attention to which one is required at each step.

#### DNA purification by magnetics beads

**Note the volume of magnetic beads used! At each stage of the protocol the amount of (NaCl+PEG) or beads and elution volume used, is stated.**

1. Add 0.65x to 2.5x sample volume of AMPure XP beads to the DNA, pipet up and down 5 times to thoroughly mix the bead suspension with the DNA, then pulse-spin and incubate at room temperature for 5 minutes.

2. Pulse-spin and place the sample tube in a magnetic rack, such as the DynaMag-2 magnet, for 3 minutes or until the solution clears. Remove and discard the supernatant without disturbing the beads.

3. Without removing the tube from the magnet, dispense 100 µL of freshly prepared 75% ethanol to the sample. Incubate for 30 seconds. After the solution clears, remove and discard the supernatant without disturbing the pellet.

4. Repeat step 3 for a second wash.

5. To remove residual ethanol, pulse-spin the tube, place it back in the magnetic rack, and carefully remove any remaining supernatant with a 20-µL pipettor without disturbing the pellet. Keeping the tube on the magnet, air-dry the beads at room temperature for 1-4 minutes until the beads are matte, but never over-dried and cracked.

6. Remove the tube from the magnet and dispense 15 or 50 µL of 1X IDTE buffer to the sample. Pipet the mixture up and down 5 times, then vortex the sample for 10 seconds, to mix thoroughly.

7. Pulse-spin and place the tube in the magnetic rack for at least 1 minute. After the solution clears, transfer the supernatant containing the eluted DNA to a new 1.5-ml Eppendorf LoBind ® tube without disturbing the pellet.

#### Beads purification when NaCl+PEG is added

**Note the volume of NaCl+PEG used! At each stage of the protocol the amount of NaCl+PEG and elution volume used, is stated.**

Same steps as above, but instead of adding AMPure XP beads, add 0.65x to 2.5x sample volume of NaCl+PEG. Pipet up and down **10 times** to thoroughly mix the NaCl+PEG with the DNA, then pulse-spin and incubate at room temperature for **10 minutes**. Then pulse-spin and place the sample tube in a magnetic rack for **5 minutes** or until the solution clears. Remove and discard the supernatant without disturbing the pellet and repeat the same steps as above.

### Appendix II: WGS Regular Coverage Calculations

For whole-genome sequencing (WGS), the required input amount for NGS-PCR vary based on the organism's genome size and the desired regular coverage depth. Use this table as a guideline for your experimental planning.

| Organism | Genome Size | Desired Regular Coverage Depth | Adapter-ligated Library Input (fmol) → Number of Total PCR Cycles | Requested Sequencing Output (Gb) |
|---|---|---|---|---|
| **Human (hg38)** | ~3 Gb | 15X | 0.033 fmol → 17 | 45 Gb |
| | | 30X | 0.066 fmol → 16 | 90 Gb |
| | | 60X | 0.133 fmol → 15 | 180 Gb |
| | | 90X | 0.2 fmol → 15 | 270 Gb |
| **Mouse (mm39)** | ~2.6 Gb | 15X | 0.029 fmol → 17 | 39 Gb |
| | | 30X | 0.058 fmol → 16 | 78 Gb |
| | | 60x | 0.116 fmol → 16 | 156 Gb |
| | | 90x | 0.175 fmol → 15 | 234 Gb |
| **Rat (GRCr8)** | ~2.8 Gb | 15X | 0.030 fmol → 17 | 42 Gb |
| | | 30X | 0.06 fmol → 16 | 84 Gb |
| | | 60x | 0.123 fmol → 16 | 168 Gb |
| | | 90x | 0.185 fmol → 15 | 252 Gb |

### Appendix III: Hybridization Input Requirements for Targeted Duplex Coverage

**TIP:** Targeted capture requires higher input DNA. We found that the use of 50 ng DNA as starting amount is enough for exome or panel. To reduce the fmol requirements during targeted capture, the use of SpeedVac is recommended during "A. Hybridization Reaction" than beads based methods.

We used the xGen™ Pan-Cancer Hybridization Panel (IDT, 1056205), which targets around 0.80 Mb of hg19 or hg38. Use the following examples to take the appropriate amount of adapter-ligated library (according to qPCR) as input for NGS-PCR and to request the appropriate sequencing output to achieve your desired duplex coverage for a sample.

| Panel | Desired Duplex Coverage (dX) | Adapter-ligated DNA to Input (fmol) | Sequencing Output to Request (Gb) |
|---|---|---|---|
| xGen™ Exome Hyb Panel v2 | ≥300 dX | 10 fmol | 60 Gb |
| xGen™ Pan-Cancer Hybridization Panel | ≥1,000 dX | 30 fmol | 60 Gb |

### Appendix IV – Examples of Expected Results

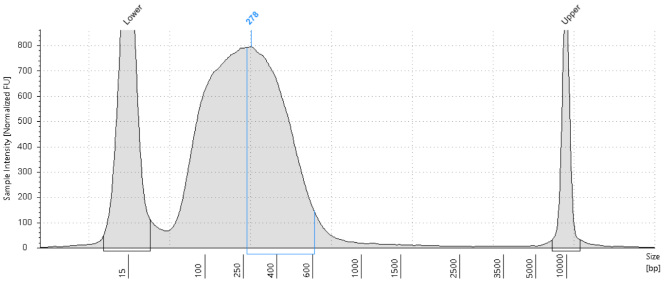

Figure A: Expected fragmented product size distribution.

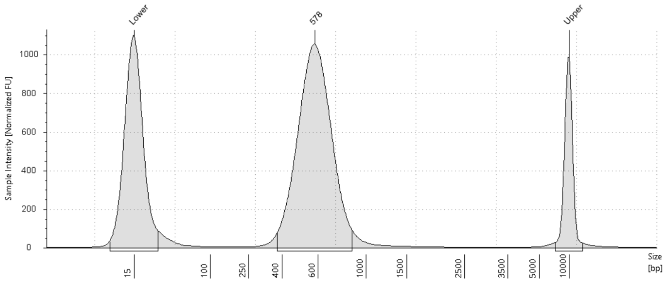

Figure B: Expected final WGS library size distribution.
